# Hybridisation and chloroplast capture between ancient *Themeda triandra* ecotypes in Australia

**DOI:** 10.1101/2021.10.21.465284

**Authors:** Luke T. Dunning, Jill K. Oloffson, Alexander S.T. Papadopulos, Paulo C. Baleeiro, Sinethemba Ntshangase, Nigel Barker, Richard W. Jobson

## Abstract

Ecotypes are distinct populations within a species which are adapted to specific environmental conditions. Understanding how these ecotypes become established, and how they interact when reunited, is fundamental to elucidating how ecological adaptations are maintained. This study focuses on *Themeda triandra*, a dominant grassland species across Asia, Africa and Australia. It is the most widespread plant in Australia, where it has distinct ecotypes that are usually restricted to either wetter and cooler coastal regions or the drier and hotter interior. We use whole genome sequencing for over 80 *Themeda* accessions to reconstruct the evolutionary history of *T*. *triandra* and related taxa. A chloroplast phylogeny confirms that Australia was colonised by *T*. *triandra* twice, with the division between ecotypes predating their arrival in Australia. The nuclear genome provides evidence of gene-flow among the ecotypes, largely restricted to two geographic areas. In northern Queensland there appears to be a hybrid zone with admixed nuclear genomes and shared plastid haplotypes. Conversely, in the cracking claypans of Western Australia, there is cytonuclear discordance with individuals possessing the coastal plastid and interior clade nuclear genomes. This chloroplast capture is potentially a result of adaptive introgression, with selection detected in the *rpoC2* gene which is associated with water use efficiency. A stable hybrid zone in the east, and the displacement of one ecotype in the west, highlights the unpredictable nature of hybrid zones, with repeated contacts between the same ecotypes producing different outcomes.

## [1] Introduction

Understanding why some species are found in a broad range of environments whilst others are restricted to particular habitats is one of the fundamental questions in evolutionary biology. Widely distributed species are typically able to tolerate multiple environmental conditions, either through extensive phenotypic plasticity or specialised ecotype differentiation (Stamp & Hadfield, 2020). It is increasingly clear that locally adapted ecotypes can evolve multiple times in response to the same selection pressures (Jones et al., 2012; Ostevik et al., 2012; Butlin et al., 2014), and that this can happen over very short timescales (Papadopulos et al., 2021). The mechanisms that a particular species utilises are dependent on numerous factors including the levels of gene flow, the extent of standing genetic variation and the scale of environmental heterogeneity (Via & Lande, 1985). Ecotypes typically arise when there is some reduced gene flow as a result of geographic isolation and strong environmental selection pressure. Determining how ecotypes interact when they come into secondary contact is important for our understanding of local adaptation and ultimately speciation (Nosil 2012).

*Themeda triandra* is one of the most widely distributed C_4_ grass species in the world (Sage, 2017). It has a relatively recent evolutionary origin, originating c. 1.5 Ma, most likely in Asia (Dunning et al., 2017). *T*. *triandra* has rapidly spread across Asia, Africa and Australia where it is found across a range of climates (Snyman et al., 2013). It is a dominant grassland species with significant ecological and economic importance (Snyman et al., 2013). It is a morphologically diverse species, and several potential taxonomic synonyms exist (Arthan et al., 2021). These synonyms range from regional varieties with restricted distributions to the globally invasive *Themeda quadrivalvis* (Arthan et al., 2021).

In Australia, *T triandra* is known to have two predominant ecotypes which are thought to be a result of ploidy differences (Godfree et al., 2017). The coastal ecotype is diploid and generally grows in cooler and wetter climates, and the inland form is prominently tetraploid and survives in drier and hotter environments (Godfree et al., 2017). Experimental manipulations show that under drought and heat stress the inland ecotype produces up to four times as much seed (Godfree et al., 2017), with elevated fitness attributed to its increased genome size. This conclusion has been supported by recent work that confirms ploidy may facilitate adaptation to hotter climates and it should be considered in adaptive management strategies (Ahrens et al., 2020). However, both of these studies based their conclusions on the assumption that the inland ecotype is derived from the coastal form, and these ploidy and ecological differences are not more ancient, predating the colonisation of Australia. Recent phylogenetic studies have highlighted that Australia may have been colonised by *T*. *triandra* multiple times (Dunning et al., 2017), but it is not yet clear how these colonisations are related to the different ecotypes.

In this study we use whole genome sequencing from over 80 *Themeda* accessions to reconstruct the evolutionary history of Themeda triandra in Australia. We specifically aim to test whether the two Australian ecotypes evolved rapidly following colonisation, potentially due to auto-polyploidisation, or alternatively, the ecotypes arrived in Australia independently with no recent auto-polyploidy event. Our results show that considering a broader evolutionary history rather than focusing purely on local diversity is essential to elucidate the mechanism of rapid environmental adaptation in widely distributed species.

## [2] Materials and Methods

### [2.1] Sampling and whole-genome sequencing

An initial Australia wide survey was conducted using an ITS/ETS barcode to assess the diversity within *T*. *triandra* / *T*. *quadrivalvis* clade. In total, we collected 332 accessions and extracted DNA, amplified the target region and generated a maximum parsimony phylogenetic tree as in Jobson et al. (2017). We then selected 64 ingroup samples distributed across this tree (Figure S1) and three outgroup samples for whole genome sequencing. The 67 selected *Themeda* accessions were then sent for sequencing at the Ramaciotti Centre for Genomics (Sydney, Australia). Libraries were constructed using the Illumina DNA Prep kit and sequenced on a single Illumina NovSeq 6000 2×150bp flow cell. Estimated genome coverage for each sample was calculated using a 1C value of 1.05 Gbp (50% of the mean tetraploid (4n) *T*. *triandra* 1C value; Estep et al., 2014).

### [2.2] Chloroplast genome assembly and phylogenetics

Chloroplast genomes were assembled from the raw whole-genome sequencing data using NOVOPlasty v.4.2.1 (Dierckxsens et al., 2017) with default parameters and a *matK* seed alignment extracted from a chloroplast genome of a closely related species (Andropogoneae: *Heteropogon* sp., GenBank accession: KY707768.1). The resulting contig options were aligned to pre-existing *T*. *triandra* and *T*. *quadrivalvis* chloroplast assemblies from NCBI genbank using MAFFT v.7017 (Katoh et al., 2002), and manually rearranged using Geneious v.5.3.6 (Kearse et al., 2012) if required. If the chloroplast assembly was incomplete, the process was repeated using a different seed alignment (the *rbcL* gene or the entire chloroplast sequence from the same *Heteropogon* sp. accession). The inverted repeat region was then removed from the alignment and a maximum-likelihood phylogenetic tree with 100 bootstrap replicates inferred using PhyML v.20120412 (Guindon et al., 2010), with the best-fit nucleotide substitution model selected with SMS v.1.8.1 (Lefort et al., 2017).

### [2.3] Nuclear marker assembly and phylogenetics

We used a reference-based approach to generate consensus sequences for single-copy nuclear genes (Olofsson et al., 2016; Dunning et al., 2019; Olofsson et al., 2019). There is no published genome for any *Themeda* species and we therefore used a non-specific reference genome from the same grass tribe (Andropogoneae: *Sorghum bicolor*, GenBank accession: GCA_000003195.3; McCormick et al., 2018). Our analysis focused on the Benchmarking Universal Single-Copy Orthologs (BUSCO; Simão et al., 2015) in the poales_odb10 database present in the *S*. *bicolor* genome. This was determined using Blastp v.2.8.1+ to identify *S*. *bicolor* proteins with a 100% sequence similarity match of at least 100 amino acids long to a sequence in the poales_odb10 database (*n* = 3,386 genes; n.b. a strict similarity cut-off was used as the poales_odb10 database includes *S*. *bicolor* proteins).

The 67 *Themeda* samples sequenced here were supplemented with 14 *T*. *triandra* and *T*. *quadrivalvis* data sets retrieved from the NCBI Sequence Read Archive (SRA; Burke et al., 2016; Dunning et al., 2017; Arthan et al., 2021). Prior to mapping the *Themeda* sequencing data to the *S*. *bicolor* genome, the data was cleaned using Trimmomatic v.0.38 (Bolger et al., 2014) to remove adaptor contamination, low quality bases (4 bp sliding window with mean Phred score < 20) and short reads (< 50 bp). NGSQC Toolkit v. 2.3.3 (Patel & Jain, 2012) was then used to discard reads where 80% of the sequence had a Phred score < 20 or the read contained an ambiguous base. Finally, PRINSEQ v.0.20.3 (Schmieder & Edwards, 2011) was used to remove duplicated reads. The cleaned data was then mapped to the *S*. *bicolor* reference genome using Bowtie2 v.2.3.4.3 (Langmead & Salzberg, 2012) with default parameters. Consensus sequences for each *Themeda* sample were then generated from the short-read alignment for each single-copy gene in the *S*. *bicolor* genome using previously described methods for low-coverage whole-genome data (Olofsson et al., 2016; Dunning et al., 2019; Olofsson et al., 2019). Each individual gene alignment was subsequently trimmed with trimAl v.1.2rev59 (Capella-Gutiérrez et al., 2009) using the -automated1 option which optimises alignment trimming for maximum-likelihood phylogenetic tree reconstruction. Short sequences (< 200 bp) were discarded from the trimmed alignment, and entire gene alignments were discarded if it was either < 500bp or did not include all samples (*n* = 1,286 genes retained).

A maximum-likelihood tree was inferred from a concatenated nuclear alignment of all 1,286 genes (alignment length = 2,532,106 bp) using IQ-TREE v.1.6.12 (Nguyen et al., 2015) with 1,000 ultrafast bootstrap replicates. Individual gene trees were also generated using SMS and PhyML as described above. A DensiTree v.2.2.7 (Bouckaert, 2010) plot was made by overlaying the individual gene trees that had been transformed to be ultrametic using the chronopl function (lambda = 1) as part of the ape v.5.2 (Paradis & Schliep, 2019) package in R v.3.4.3. A coalescence species tree was generated from the individual gene trees using ASTRAL v.5.7.5 (Zhang et al., 2018) after collapsing branches with <10% bootstrap support using Newick utilities v.1.6 (Junier & Zdobnov, 2010). Phyparts v.0.0.1 (Smith et al., 2015) was used to evaluate individual gene tree support for the coalescence species tree. The results were visualised using the phypartspiecharts.py python script written by M. Johnson (available from: https://github.com/mossmatters/phyloscripts/blob/master/phypartspiecharts).

### [2.3] Population structure

Genotype likelihoods were estimated across the nuclear genome of the 81 *Themeda* acessions (67 sequenced here) using ANGSD v.0.929-13-gb5c4df3 (Korneliussen et al., 2014) and the bowtie2 alignments generated above. Mapped reads and bases with a quality < 20 were discarded, a per-individual maximum depth of 20 was used, and sites had to be present in at least two individuals to be considered. A principal component analysis (PCA) of the genotype likelihoods was generated using PCAngsd v.0.973 (Meisner & Albrechtsen, 2018) to estimate a covariance matrix before plotting the results in R with eigenvector decomposition. The number of genetic clusters (K) in the genotype likelihoods was examined using NGSadmix (Skotte et al., 2013), with default parameters and 10 replicates for K between 1 and 10. The optimal K was determined using the ΔK method (Evanno et al., 2005) implemented using CLUMPAK (Kopelman et al., 2015). PCA plots and the admixture analysis were repeated for only the Australian *T*. *triandra* samples using default parameters. Pairwise *F*_ST_ was estimated using ANGSD among Australian populations restricted to individuals that had 99% of the genomes assigned to a single genetic cluster both globally and using a sliding window method (window = 50kb, slide = 10kb).

### [2.4] Estimating ploidy

The ploidy of each sample was estimated using HMMploidy (Soraggi et al., 2021), a method which has been developed to infer ploidy from low-depth sequencing data. A multi-sample mpileup file was generated for HMMploidy from the bowtie2 alignments with SAMtools v.1.9 (Li et al., 2009), only including reads with a minimum read mapping quality (mapQ) of 20, counting anomalous read pairs and setting a maximum per-file depth of 25. Genotype likelihoods were generated using HMMploidy with default parameters, which calculates likelihoods for a range of ploidy levels (1n–6n). Ploidy levels were then inferred in 100kb windows across the *S*. *bicolor* reference genome, with a minimum number of two individuals per locus to be considered. The percentage of 100kb windows supporting each ploidy level was then calculated, ignoring those that were inferred to be haploid. Samples were arbitrarily assigned to a single ploidy if it was supported by over 50% of windows, or ‘likely’ a certain ploidy if only one category was supported by over 40% of windows. If samples were not assigned to a single ploidy level, but over 60% of windows supported a ploidy level greater than diploid, then they were classified as ‘likely polyploid’, or ‘unknown’ if less than 60%.

### [2.4] Inferring positive selection

Pairwise estimation of the ratio (ω) of synonymous (dS) and non-synonymous (dN) substitutions between sequences was calculated using the Yang and Nielson (2000) method implemented in yn00, distributed as part of the paml v.4.9j package (Yang, 2007). Site (M1a and M2a) and branch-site models (BSA and BSA1) models to infer if a gene was evolving under significant positive selection were implemented in codeml, distributed as part of the paml v.4.9j package (Yang, 2007). For these models we used the topology of the whole chloroplast genome tree (Figure 1 & S1).

**Figure 1:**
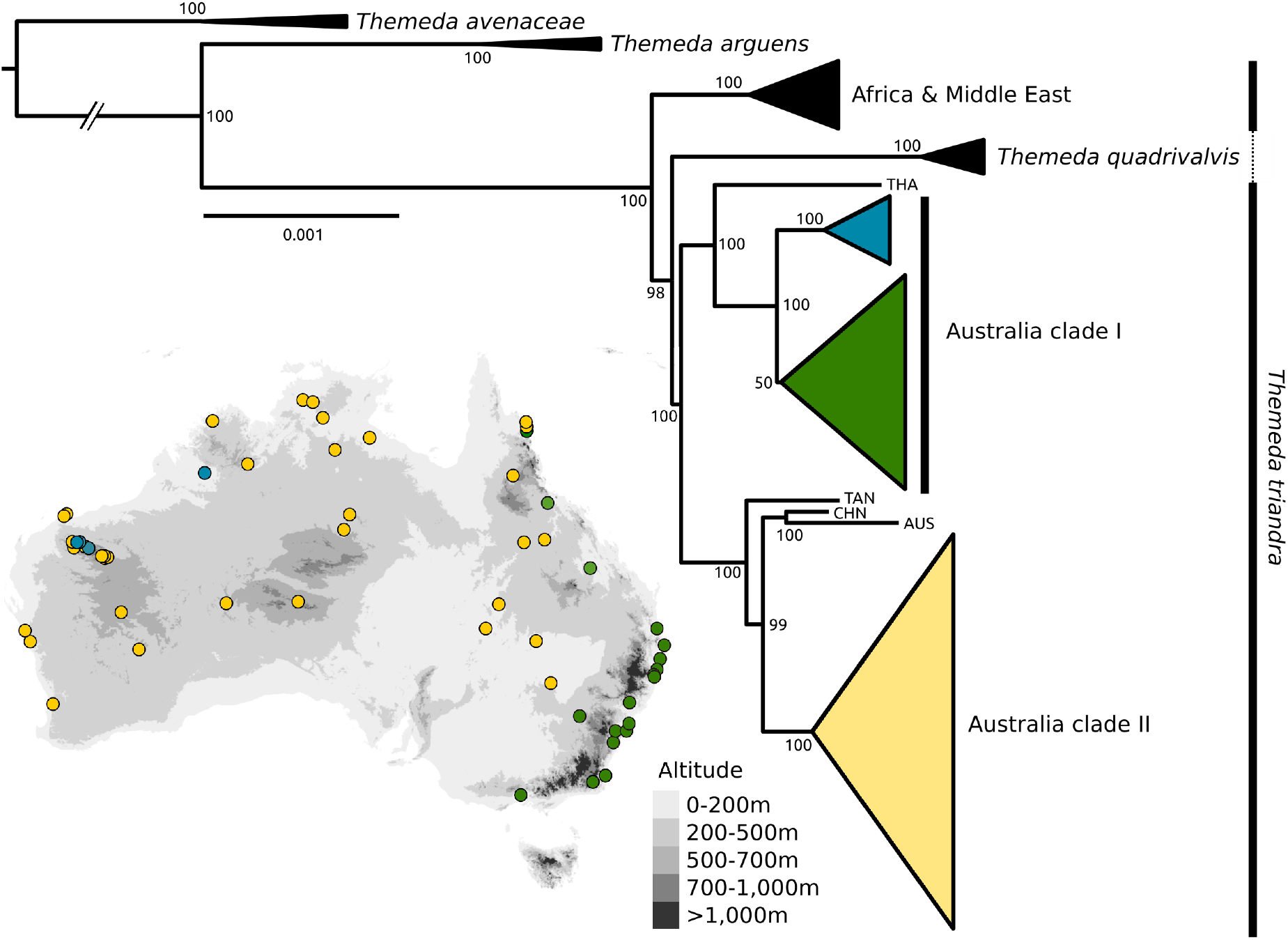
Phylogenetic relationships of *Themeda* inferred from whole chloroplast genomes. The maximum likelihood topology inferred with the GTR+I+G substitution model is shown with bootstrap support values. The sampling location of *T*. *triandra* accession in Australia Clade I and II is shown, and truncated branches are indicated. For samples not assigned to a clade a three letter abbreviation is used (THA = Thailand; TAN = Tanzania; CHN = China; AUS = Australia).

## [3] Results

### [3.1] Phylogenetic relationships inferred from the chloroplast

As part of this study we generated over 600 Gb (mean = 4.6 Gb, SD = 1.4 Gb per sample) of whole-genome sequencing data for 67 *Themeda* accessions, with an emphasis on sampling *T*. *triandra* (and potential taxonomic synonyms) from across its range in Australia (*n* = 61; Figure 1 & S2). These data were used to assemble whole chloroplast genomes, before being aligned with previously published assemblies (*n* = 15). Inferring the phylogenetic relationships based on the chloroplast genomes (82 accessions in total, 118,550 bp alignment, 95.4% identical sites, 97.9% within *T*. *triandra*; 98.6% and 99.1% when excluding indels) shows that *T*. *triandra* is not monophyletic, with *T*. *quadrivalvis* nested within. The phylogeny also shows that the Australian samples do not form a single clade (Figure 1), as would be expected if there was a single colonisation of Australia. The two large Australian clades (Clade I & II) are well supported, each sister to Asian accessions. Clade I comprises the coastal *T*. *triandra* accessions predominantly from wetter environments (in green) and those from the Pilbara cracking claypans in Western Australia (in blue; Figure 1). Clade II is composed of the western *T*. *triandra* form (in yellow), which are predominantly from dryer habitats in the Australian interior. When comparing the coastal (green) and claypan (blue) *T*. *triandra* accessions from Clade I with the western Australia form from Clade II (yellow), there are 48 and 64 fixed biallelic SNPs respectively. Within Clade I there are 13 fixed biallelic SNPs separating the coastal (green) and Pilbara (blue) *T*. *triandra* accessions.

### [3.2] Phylogenetic relationships inferred from the nuclear genome

The nuclear dataset was generated by mapping the short-read data from 81 *Themeda* accessions to the *S*. *bicolor* genome. Using a non-specific reference reduces the proportion of data that maps (mean = 10.64%, SD = 1.35%), with valid alignments mainly restricted to exons (Olofsson et al., 2019). The nuclear relationships were inferred using data from 1,286 single-copy genes with a combined alignment length of 2,532,106 bp. The concatenated maximum-likelihood tree and the coalescent species tree both support *T*. *triandra* as being monophyletic, sister to *T*. *quadrivalvis* (Figure 2, S3–S5). In the nuclear phylogeny the Australian *T*. *triandra* accessions are monophyletic, and within this group the two distinct clades identified in the chloroplast phylogeny are recapitulated with a few exceptions. Most notably is the Pilbara clade which is identified as Clade I based on the chloroplast (in blue; Figure 1), but is nested deep within Clade II based on the nuclear genome (Figure 2). There are also two accessions from Far North Queensland (FNQ) which are part of Clade II in the chloroplast genome which are nested within Clade I based on the nuclear genome, where they form a clade with other accessions from FNQ. Evaluating the individual gene tree support for the nuclear topology shows that many of the nodes are poorly supported beyond delimiting the main clades (Fig 2 & S5). This lack of gene-tree support for the species tree topology can be a result of incomplete lineage sorting, hybridisation and/or a lack of genetic variation.

**Figure 2:**
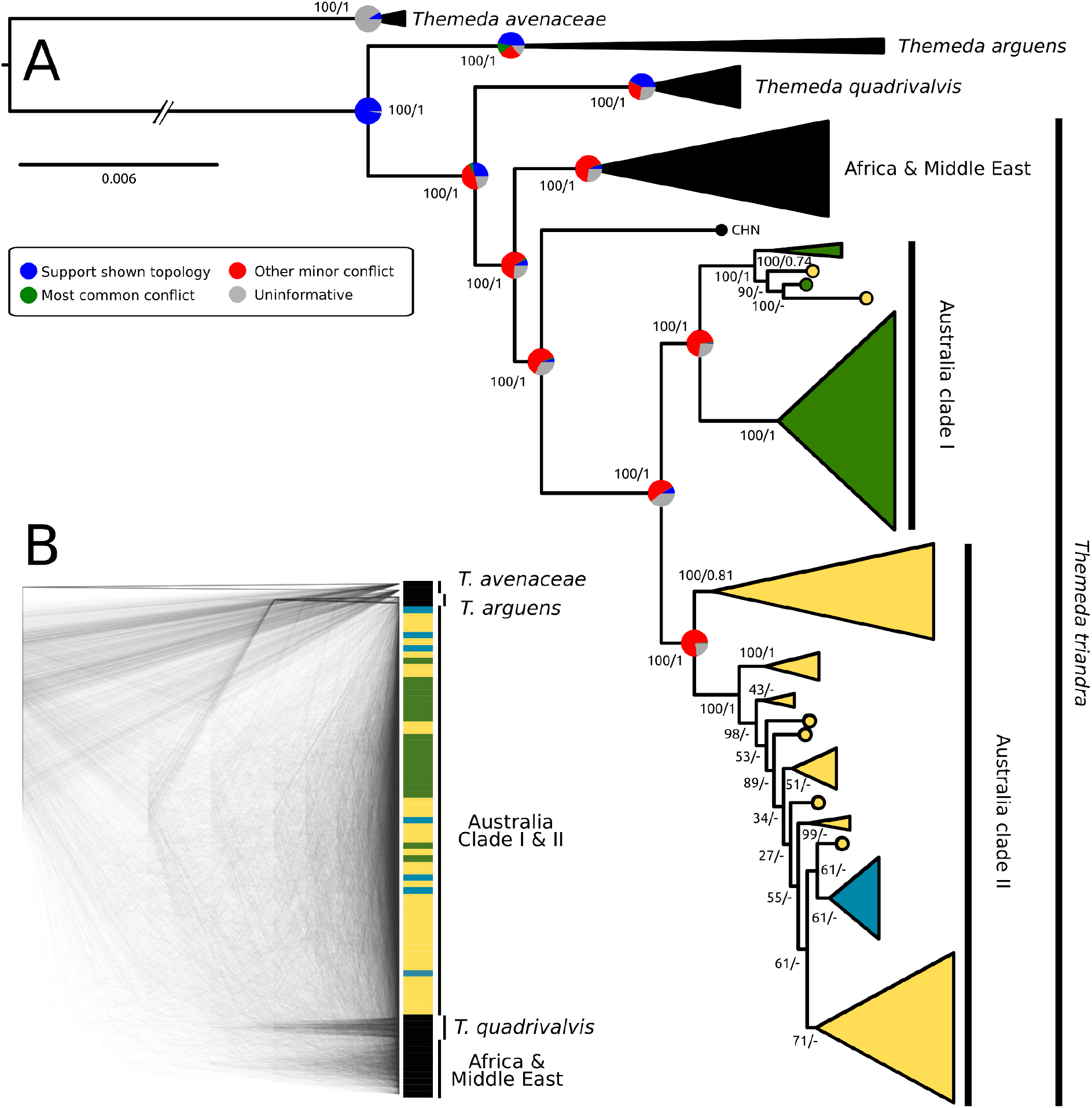
Phylogenetic relationships of *Themeda* inferred from 1,286 nuclear genes. A shows a maximum likelihood topology from a concatenated alignment of all loci with colours of the Australian clades based on chloroplast groupings (Figure 1). Bootstrap support values for the concatenated maximum likelihood tree are shown, followed by local posterior probabilities from a coalescence species tree for the same nuclear loci. The pie charts on key nodes represent the individual gene tree support for the topology shown. B shows a densitree plot of overlaid nuclear gene trees. Colours match the chloroplast clades, and truncated branches are indicated.

### [3.3] Genetic variations and structure within *Themeda*

The principle component analysis largely recovered the nuclear phylogeny groupings (Figure 3A). The first principal component axis explains 27.0% of the variation in the data and predominantly splits the *T*. *triandra* samples from the other accessions, although they themselves are clustered in their nuclear phylogeny clades. The second principal component explains 16% of the variation in the data and splits the Australian accession into the two distinct nuclear clades, with a small group of five accessions from FNQ and the surrounding area. The dominance of Australian *T*. *triandra* accessions in the PCA is likely to be a direct result of the high proportion (79.0%) of samples these accessions comprise. We therefore repeated the analysis, only including Australian *T*. *triandra* accessions, with similar results (Figure S6).

**Figure 3:**
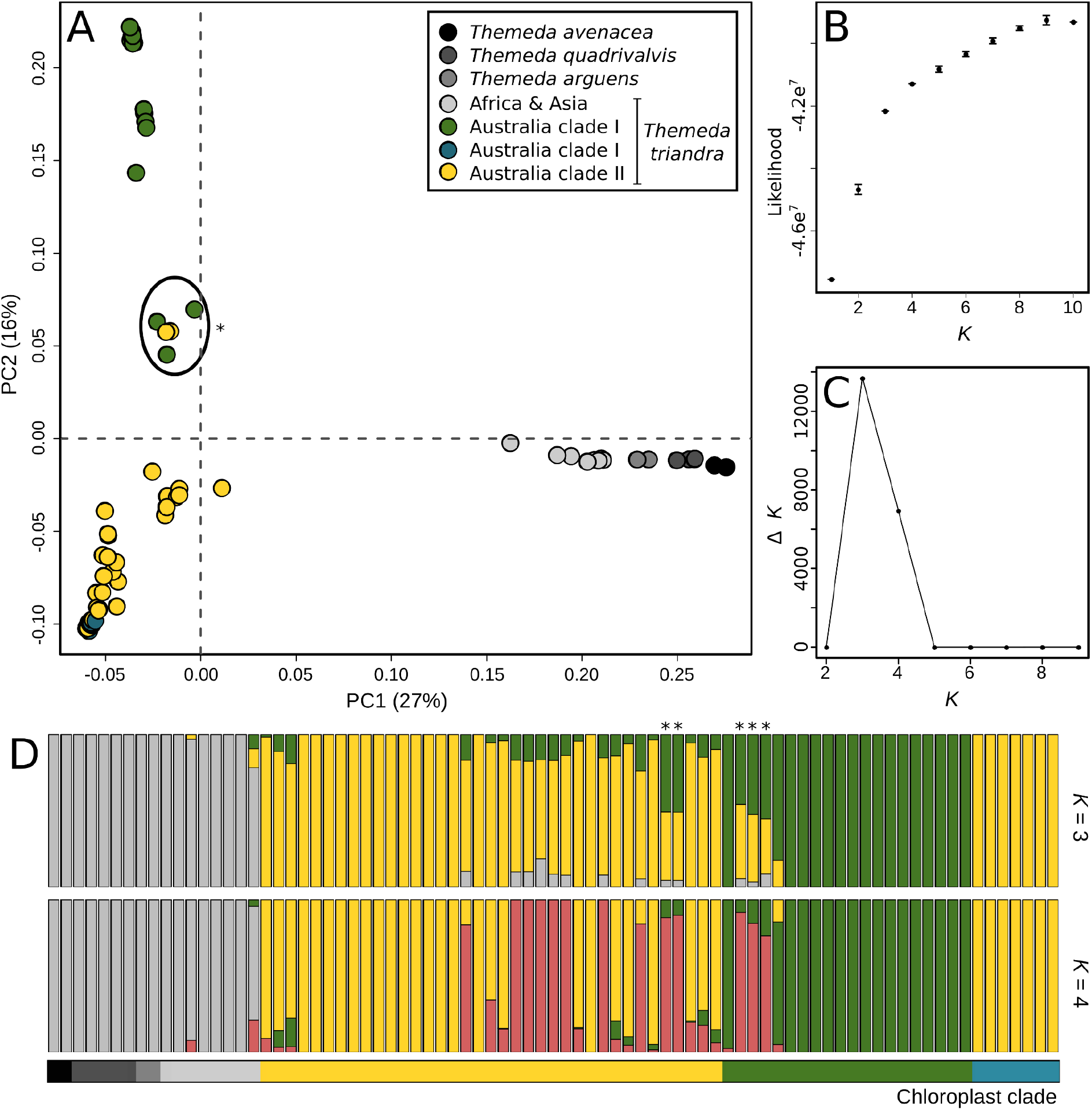
Genetic variation and structure within *Themeda*. (A) A principle component analysis across the first two axes is shown, with genetic groups coloured based on the chloroplast phylogeny as in Figure 1. (B) The mean likelihood and standard error for a range of *K*’s is shown, with these values used to calculate Δ*K* (C) as in Evanno et al., (2005). The assignment to genetic clusters is shown for two values of *K*. Samples are arranged within their chloroplast clade (indicated by that bar underneath the admixture plots), and ordered from west to east within each group. The asterisks indicate samples with a high degree of admixture.

The optimal number of genetic clusters (*K*) based on the admixture analysis is three, with a secondary optimum of four (Fig 3B & C). For both *K* = 3 and *K* = 4 the accessions outside of the Australian *T*. *triandra* are lumped into a single isolated group, although the *T*. *triandra* accession from China has between 10 and 20% of its genome assigned to one of the other genetic clusters (Figure 3D). The most optimal *K* (3) largely recovers the nuclear phylogeny, confirming that the Pilbara claypan accessions have a mis-match between their chloroplast (Clade I) and nuclear (Clade II) genomes. The samples are ordered from west to east in each chloroplast genotype block, indicating that there is increased admixture where the two clades come into contact (Figure 3D). There is an extremely high-degree of admixture in five individuals with roughly equal proportions of their nuclear genome assigned to Clade I and Clade II (Figure 3D). These are the same five individuals from FNQ and the surrounding area which were intermediate in the PCA (Figure 3A). These individuals also make up the small nuclear clade with mixed chloroplast genotypes in Figure 2. The secondary optimum *K* = 4 recovers the same pattern, although it largely assigns individuals with visible admixture in *K* = 3 to an additional 4th genetic group (Figure 3D). The same groupings are largely recaptured when repeating the PCA and admixture analysis (optimum *K* = 3; secondary optimum *K* = 2) with only Australian *T*. *triandra samples* (Figure S5). With this reduced sample set PC1 accounts for 26% of variation and has a significant correlation with sample longitude (Pearson’s r = 0.84; P-value < 0.001).

Pairwise *F*_ST_ confirmed that the nuclear genome of the claypan form and inland ecotype are more similar (*F*_ST_ unweighted = 0.06; *F*_ST_ weighted = 0.19) to each other than to the coastal ecotype (*F*_ST_ unweighted = 0.22; *F*_ST_ weighted = 0.72; and *F*_ST_ unweighted = 0.18; *F*_ST_ weighted = 0.70, respectively). Pairwise *F*_ST_ across the genome showed a similar pattern (Figure S7).

### [3.4] Polyploidy is restricted to the inland ecotype

All outgroups and accessions of the Australian coastal ecotype were assigned as diploid, or likely diploid (Table S2). Accessions from the interior ecotype were inferred to have a range of ploidies ranging from diploid to hexaploid, although a majority (66%) of samples were polyploids (Table S2). No sample from this clade was assigned as tetraploid, even though sampling in southeastern Australia indicated this is the most common ploidy for the inland ecotype (Godfree et al., 2017; Ahrens et al., 2020). It is likely that in our analysis tetraploid individuals were assigned to the ‘likely polyploid’ category in our results. The introgressed individuals of both ecotypes from FNQ were diploid, indicating that these were not allopolyploids and likely represent early-generation hybrids. The ploidy assignments were largely consistent across the different chromosomes, with some differences potentially a result of aneuploidy which has been documented in Australian individuals (Hayman, 1960).

### [3.5] Positive selection in a single chloroplast gene

We looked at sequence variation within all 75 protein coding chloroplast genes to detect signs of positive selection between the two Australian *T*. *triandra* clades potentially involved in a chloroplast capture scenario (i.e. blue and yellow clades; Figure 1). Out of the 75 genes, 60 had no fixed differences, 11 had fixed differences but they were all synonymous mutations, and four (*ndhf*, *rpl22*, *rpoA*, and *rpoC2*) contained non-synonymous mutations. Only *rpoC2* had more than three fixed differences and was therefore used for positive selection analysis. A simple pairwise comparison between sequences using the Yang and Nielsen (2000) method indicates *rpoC2* had been under positive selection between the Australian clades, with an excess of non-synonymous mutations (ω = 1.39; dN = 0.0012; dS = 0.0009). This conclusion is supported when testing for positive selection using more complex site models across the phylogeny (M2a > M1a; *P*-value < 0.001), which identified one site with >95% probability as being under positive selection (site 1503; P = 98.9%). The amino acid residue at this site is divergent between the clades of interest as a result of a non-synonymous mutation that has arisen on the branch separating the Chinese/Australian Clade II samples from the Tanzanian accession (Figure 1). However, branch-site models did not find any significant evidence for elevated positive selection on the branches separating the two clades of interest compared to the rest of the tree (Table S3).

## [4] Discussion

*Themeda triandra* is one of the most charismatic tropical grass species. Despite being relatively young (Dunning et al., 2017), this species has become dominant in many African, Asian and Australian grasslands, and has even been dubbed the ‘The food of the Serengeti grazers’ (Sage, 2017). It is the most widely distributed plant species in Australia (Gallagher 2016) with two predominant ecotypes largely restricted to the wetter, cooler coastal regions or the drier, hotter interior. It is therefore a great model system to investigate rapid environmental adaptation. Here, we use whole-genome sequencing from over 80 *Themeda* accessions to show that divergence between the ecotypes pre-dates the colonisation of Australia by *T*. *triandra*, and they are therefore not a result of a recent auto-polyploidisation event. We also show extensive gene-flow between ecotypes with varying outcomes.

### [4.1] Ecotypes predate the colonisation of Australia

This study confirms previous findings that Australia was colonised at least twice by *Themeda triandra* (Dunning et al., 2017). Interestingly, these independent colonisations actually represent the arrival of the two different ecotypes, one that inhabits the cooler and wetter coastal regions and the other that is found in the hotter and drier Australian interior (Figure 1). Previous studies have concluded that the differences between these ecotypes can be attributed to recent polyploidy events (Godfree et al., 2017; Ahrens et al., 2020). While our results support these previous conclusions that ecotypes do largely segregate by ploidy level (Table S2), the adaptation to these different environments likely pre-dates the colonisation of Australia and is not solely the result of a recent auto-polyploidisation event. Indeed, we detect diploid individuals within the dryland interior ecotype. Therefore, to understand the colonisation of the dryland Australian interior it is essential to consider the ecotype’s original diversification in Asia. A recent study of *T*. *triandra* in Yunnan-Guizhou Plateau in southwest China also found distinct cool- and warm-adapted lineages that are at least 2 million years old (Chu et al., 2021). However, the distinct Chinese populations are unlikely to be the source of the two Australian ecotypes as the Chinese samples form a monophyletic group based on chloroplast markers, although this is based on a reduced set of markers and a topology with relatively low support (Chu et al., 2021).

*Themeda triandra* is also widespread in Africa, and grows in a wide range of climatic regions and exhibits similar diversity in ploidy levels (Snyman et al., 2013). Although presently not possible, it would be interesting to compare the Australian results with a similar study in Africa. Potentially the African continent was also colonised by multiple ecotypes, and indeed African accessions have a similar phylogenetic pattern being paraphyletic for the chloroplast genome (placement of Tanzanian accession, Fig. 1) and monophyletic for the nuclear genome (Fig. 2). Comparison between Australia and Africa will ultimately show if the broad climatic niche *T*. *triandra* inhabits on both continents is attributed to ancestral genetic variation or rapid convergent evolution.

### [4.2] Hybridisation between ecotypes

The nuclear phylogenies and population genomics results indicate that there is ongoing gene flow between the ecotypes where they come into contact (Figure 2 & 3). The highest levels of introgression is localised to Far North Queensland (FNQ) where diploid accessions of both ecotypes are found in close proximity and which have the appearance of early generation hybrids (Figure 2). Populations at increasing distance from this potential hybrid zone contain successively reduced introgression to populations with a relatively pure nuclear background for each ecotype. *Themeda triandra* only relatively recently colonised Australia (< 1.3 Mya; Dunning et al., 2017) and at present it is unclear how stable this hybrid zone in Far North Queensland is, and it may represent a profitable geographic location to investigate the genetic basis of the two ecotypes. It is also likely that the predominant ploidy differences between ecotypes also aids in maintaining the divergence between ecotypes (Olofsson et al., 2021)

There is also potentially ongoing gene-flow from Asia into northern Australia. When including all accessions in the structure analysis the Chinese sample shares noticeably high levels of population assignment to the Australian samples from northern Australia, and at k=4 these regions separate into their own 4th cluster (Figure 3). This might also explain why when restricting the analysis to only Australian accessions k=3 remains the optimal number of populations as it is still detecting the signal of introgression into these accessions. Further sampling, particularly in Southeast Asia, is required to confirm this conclusion.

### [4.3] Chloroplast capture in cracking claypans of Western Australia

The Western Australian cracking claypans around Pilbara are characterised by frequent inundation with fresh water, compared to the surrounding drier desert regions. *Themeda triandra* accessions in these restricted habitats have been previously classified as a separate species (*Themeda* sp. Hamersley Station) based on morphological differences, although subsequent inspection by taxonomists have shown these differences are largely qualitative. The genetic data indicate that populations from these areas are in fact *Themeda triandra* (Figure 1 & 2). However, there is clear nuclear & chloroplast discordance in these accessions, possessing an interior ecotype nuclear genome and a coastal ecotype chloroplast. A comparison of the coding genes in the chloroplast indicates that one gene (*rpoC2*) in particular has an ω > 1 indicating positive selection between ecotypes. This may mean that the observed chloroplast capture was a result of adaptive introgression.

The *rpoC2* gene encodes a DNA-dependent RNA polymerase, and its expression has been previously associated with increased water use efficiency in the common bean (*Phaseolus vulgaris*; Ruiz-Nieto et al., 2015). Comparative analysis of rice chloroplast genomes have also shown *rpoC2* to be under positive selection in *Oryza* species from high light environments, and in particular *Oryza autraliensis*, a wild rice native to northern Australia (Gao et al., 2019). However, when attempting to determine if the amino acid substitutions are convergent between *O*. *autraliensis* and *T*. *triandra* it became clear that the positive selection result in the former are a likely a false-positive driven by poorly aligned indel regions rather than actual amino acid substitutions. The role of *rpoC2* in water use efficiency, and the difference in water availability in the cracking claypans versus the surrounding habitat, potentially indicates that adaptive chloroplast capture of the coastal chloroplast has occurred. This pattern might also provide support for a once widespread coastal ecotype across Australia, including its interior, which has then been largely replaced by the interior ecotype as the continent oscillated in water availability before trending to become drier in the last 350,000 years (Kershaw et al., 2003).

### [4.4] Recent speciation of *Themeda quadrivalvis*

Whether *Themeda quadrivalvis* is a synonym of *Themeda triandra*, or if it is a separate species has been debated for some time (Keir & Vogler, 2006; Veldkamp et al., 2016; Dunning et al., 2017; Arthran et al., 2021). *Themeda quadrivalvis* is a globally distributed invasive weed and the only apparent fixed difference between species is that *T*. *quadrivalvis* is annual whereas *T*. *triandra* is perennial. This is the first study to sequence multiple genomes of *T*. *quadrivalvis* and the results support a previous conclusion that this species has only recently diverged from *T*. *triandra*. In the early stages of speciation, a daughter species would sit within the larger paraphyletic parental species (Pennington & Lavin 2016). This is exactly what we observe in the slower evolving chloroplast genome (Figure 1), whereas each species is monophyletic in the nuclear genome (Figure 2). Despite occurring in the same geographic location within Australia, we fail to detect any meaningful gene flow between these putative species (Figure 3), suggesting that they are now largely reproductively isolated.

### [4.5] Conclusion

*Themeda triandra* represents one of the most recent and successful rapid radiations of grasses. In a relatively short space of time, it has become the most broadly distributed plant species within Australia across a very broad ecological spectrum. This remarkable feat is likely due to ecological plasticity. However, this research shows that this ability to occupy almost every climatic niche in Australia is a result of independent colonisation of the continent by ecotypes within this species, with some further local adaptation potential arising following introgression of chloroplast haplotypes. This results highlights the importance of standing genetic variation within a species in facilitating rapid adaptation to a diverse array of habitats.

## [5] Acknowledgements

We thank Natalie Murdock and Hayden Ajduk (Rio Tinto Mining), Steven Dillon (PERTH), and Andrew Mitchell (Perth, WA) for help with images, fieldwork and identification of specimens. We are grateful for access to collections and provision of loans by managers and staff at AD, AK, BRI, CANB, CNS, DNA, HO, MEL, NSW, PERTH, SING. The Australian Biological Resource Study (ABRS) and Rio Tinto Mining provided funding for this research. LTD is supported by a Natural Environment Research Council Independent Research Fellowship (NE/T011025/1).

## [7] Data Accessibility

All raw whole-genome sequencing data has been deposited in the NCBI Sequence Read Archive (Bioproject XXXX). Chloroplast genomes were submitted to NCBI GenBank (Accession numbers XXXX-XXXX).

## [8] Author Contributions

L.T.D and R.J conceived and designed the research with contributions from all coauthors. P.C.B, S.N, N.B and R.J conducted fieldwork and generated initial ITS/ETS data. L.T.D, J.K.O and A.S.T.P analysed the whole-genome data. L.T.D. and R.J. wrote the manuscript and all authors commented on the final version.

## [9] Supplementary Tables and Figures

**Table S1:** Sample details:

Available from here:

https://docs.google.com/spreadsheets/d/1UcJz1HDWF8r2hDwboRzetbAbp_SG0ZUm/edit?usp=sharing&ouid=114024510845369830312&rtpof=true&sd=true

**Table S2:** Inferring ploidy level results. Samples are ordered as they are in the nuclear phylogeny (Figure S3):

Available from here:

https://docs.google.com/spreadsheets/d/1BzZySUNc3SBSi-MXWo0uSinBeRS6GhjR/edit?usp=sharing&ouid=114024510845369830312&rtpof=true&sd=true

**Table S3:** Results of the branch-site models used to detect positive selection. See figure S2 for the topology referred to.

| Model | Likelihood | P-value | Daughter Clade |
| --- | --- | --- | --- |
| Null model | -6772.90 | - | - |
| Branch 1 | -6772.90 | 1.00E+00 | Claypan clade |
| Branch 2 | -6772.23 | 2.49E-01 | Australian Clade I |
| Branch 3 | -6772.90 | 1.00E+00 | Australian Clade I + Thailand |
| Branch 4 | -6772.22 | 2.45E-01 | Australian Clade II |
| Branch 5 | -6772.90 | 1.00E+00 | Australian Clade II + China |
| Branch 6 | -6772.26 | 2.57E-01 | Australian Clade II + China + Tanzania |

**Figure S1:**
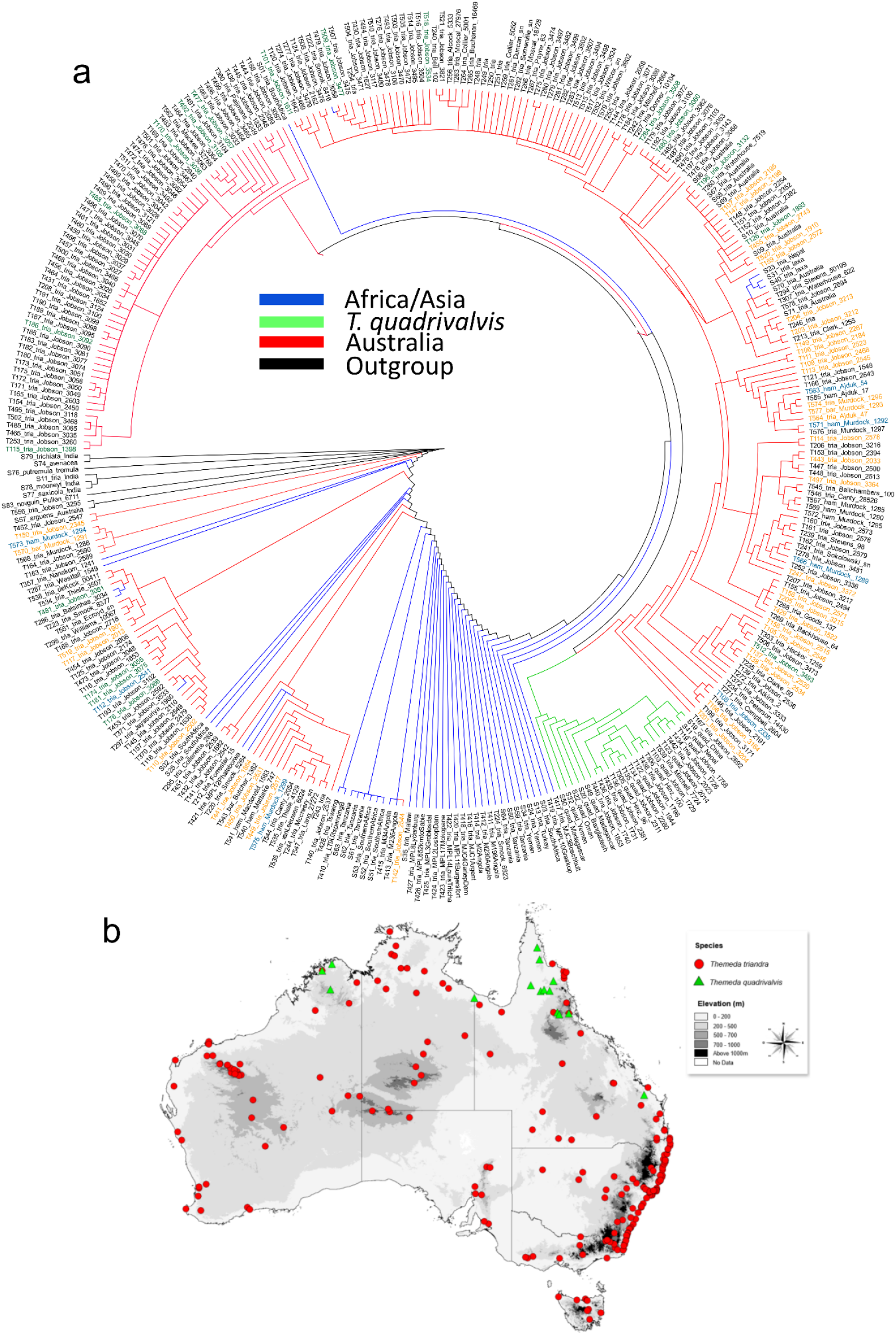
Initial survey of *Themeda* diversity in Australia. (a) A maximum parsimony topology is inferred from the ITS/ETS barcode region for 332 accession. A subset of 67 accessions were subsequently selected for whole genome resequencing to ensure the diversity of Australian genotypes was sampled. (b) the geographic distribution of Australian samples is shown.

**Figure S2:**
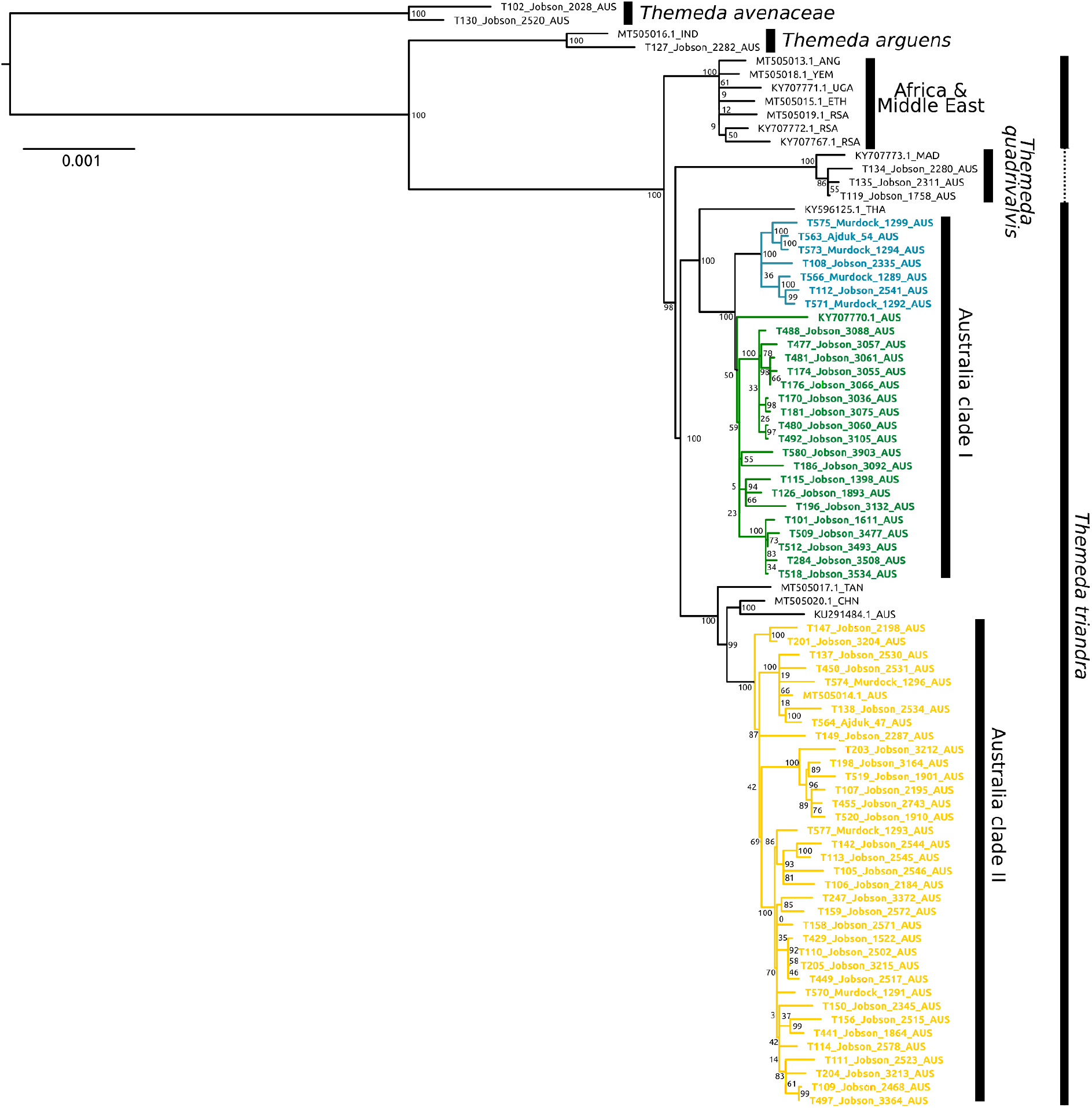
Phylogenetic relationships of *Themeda* inferred from whole chloroplast genomes. The maximum likelihood topology is shown with bootstrap support values based on 100 bootstrap replicates

**Figure S3:**
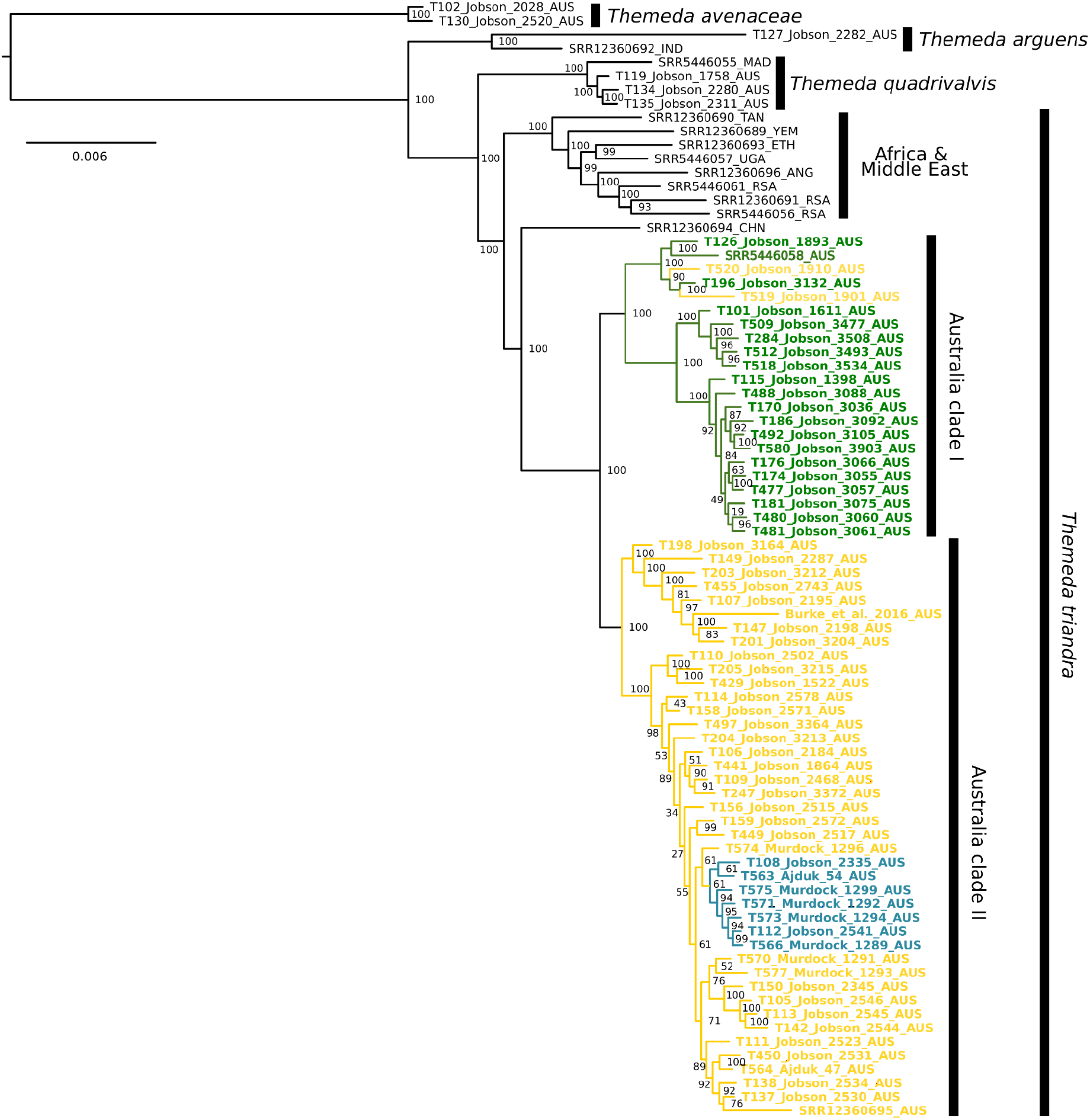
Phylogenetic relationships of *Themeda* inferred from a concatenated nuclear alignment of 1,286 genes. The maximum likelihood topology is shown with bootstrap support values based on 1,000 rapid bootstrap replicates. Colours are based on chloroplast clade.

**Figure S4:**
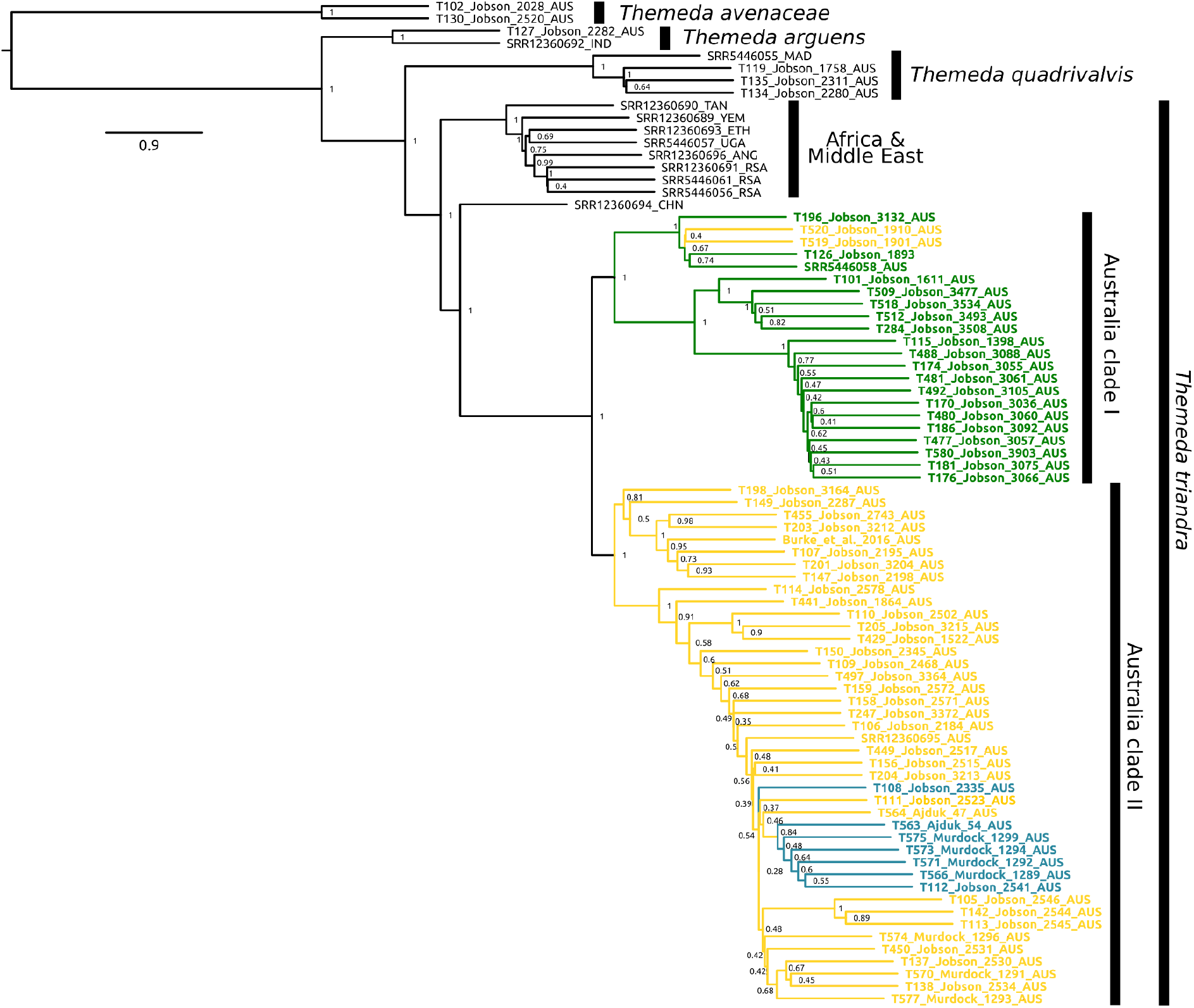
Phylogenetic relationships of *Themeda* inferred from 1,286 genes. The coalescence species tree is shown with local posterior probabilities. Branch lengths are in coalescence units, arbitrarily set to one for the tips. Colours are based on chloroplast clade.

**Figure S5:**
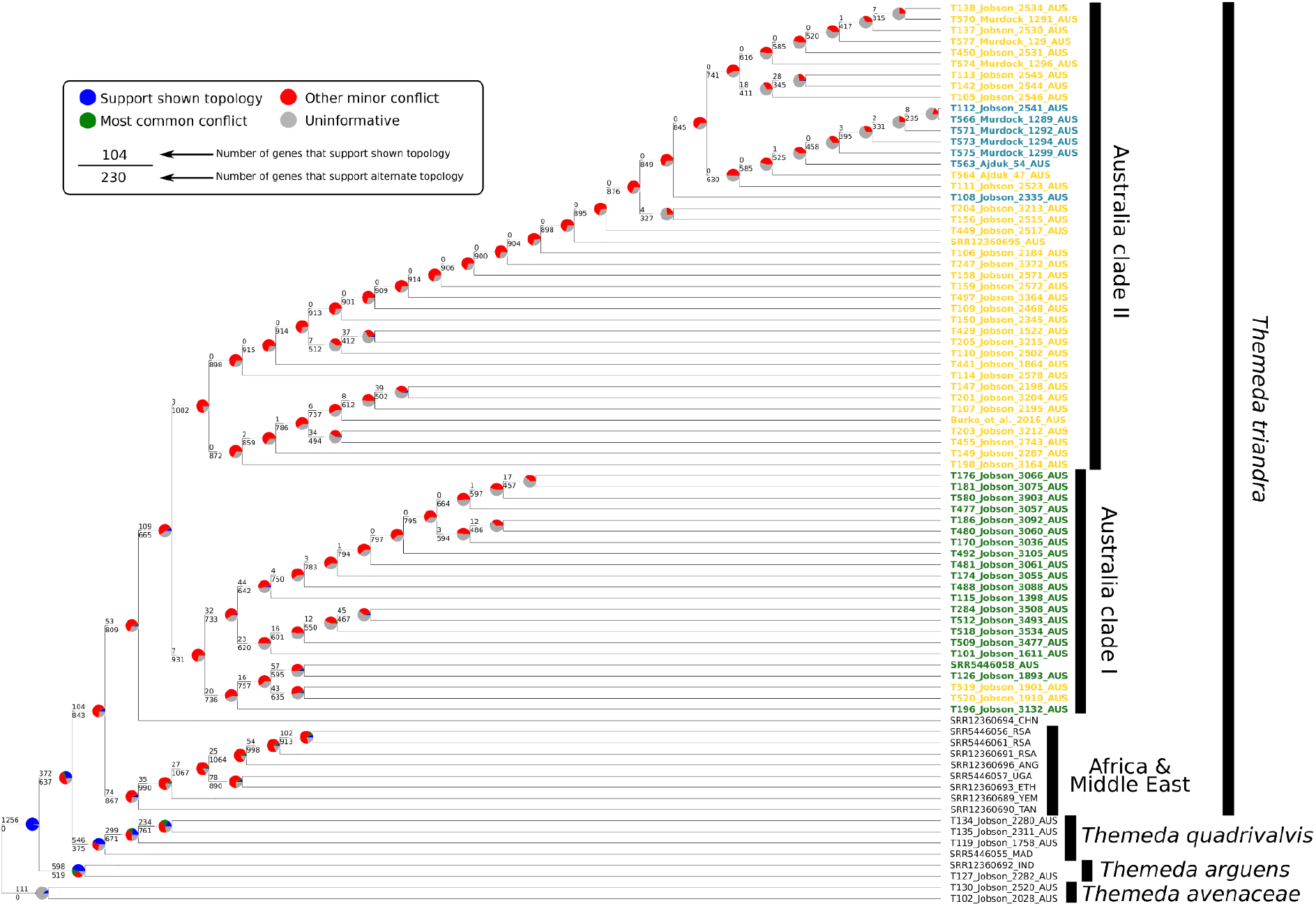
Evaluating individual gene tree support for the coalescence species tree topology. Branch lengths are arbitrary. Colours are based on chloroplast clade.

**Figure S6:**
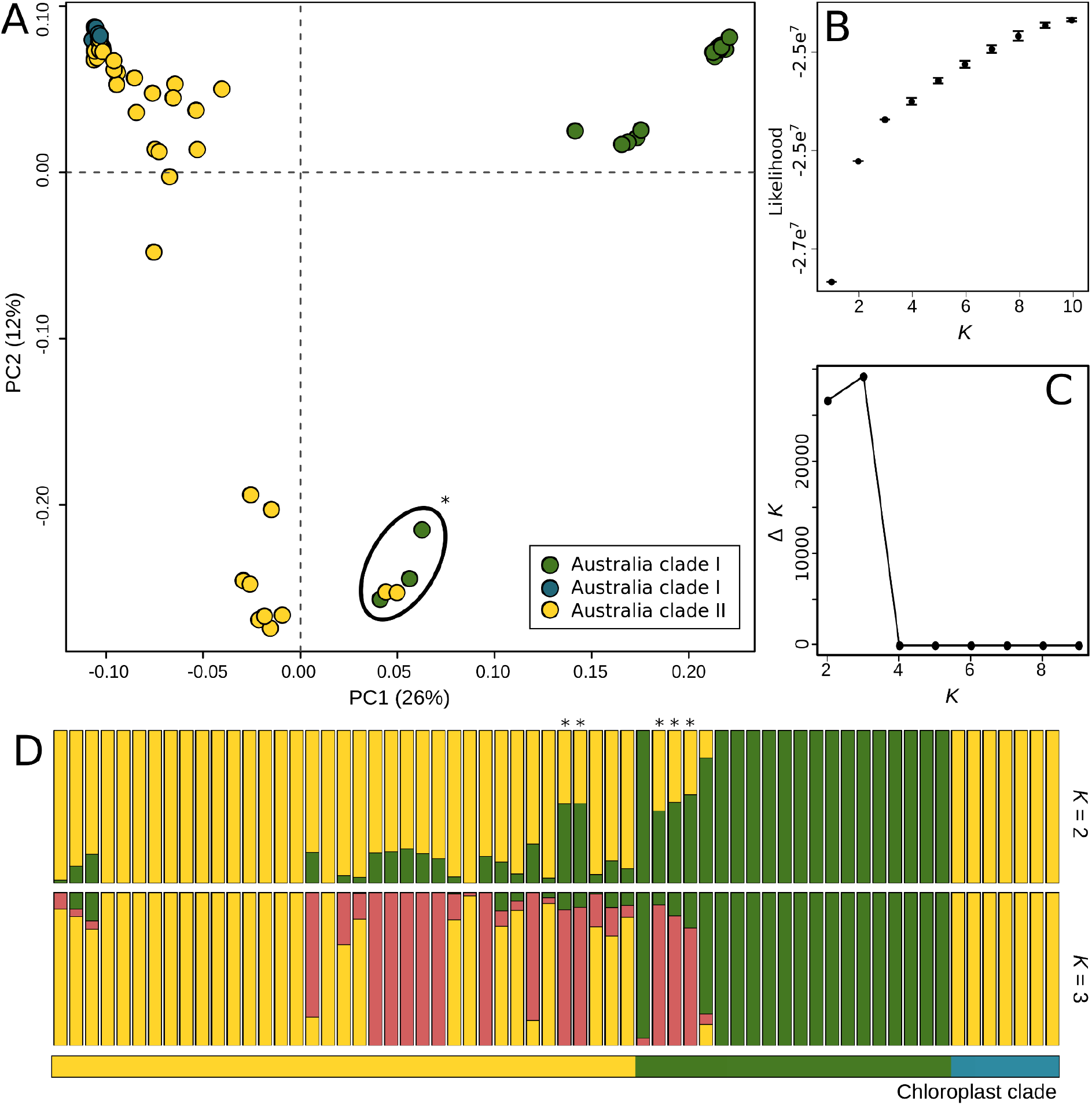
Genetic variation and structure within *Themeda triandra* in Australia. (A) A principle component analysis across the first two axes is shown, with genetic groups coloured based on the chloroplast phylogeny as in Figure 1. (B) The mean likelihood and standard error for a range of *K*’s is shown, with these values used to calculate Δ*K* (C) as in Evanno et al., (2005). The assignment to genetic clusters is shown for two values of *K*. Samples are arranged within their chloroplast clade (indicated by that bar underneath the admixture plots), and ordered from west to east within each group. The asterisks indicate samples with a high degree of admixture.

**Figure S7:**
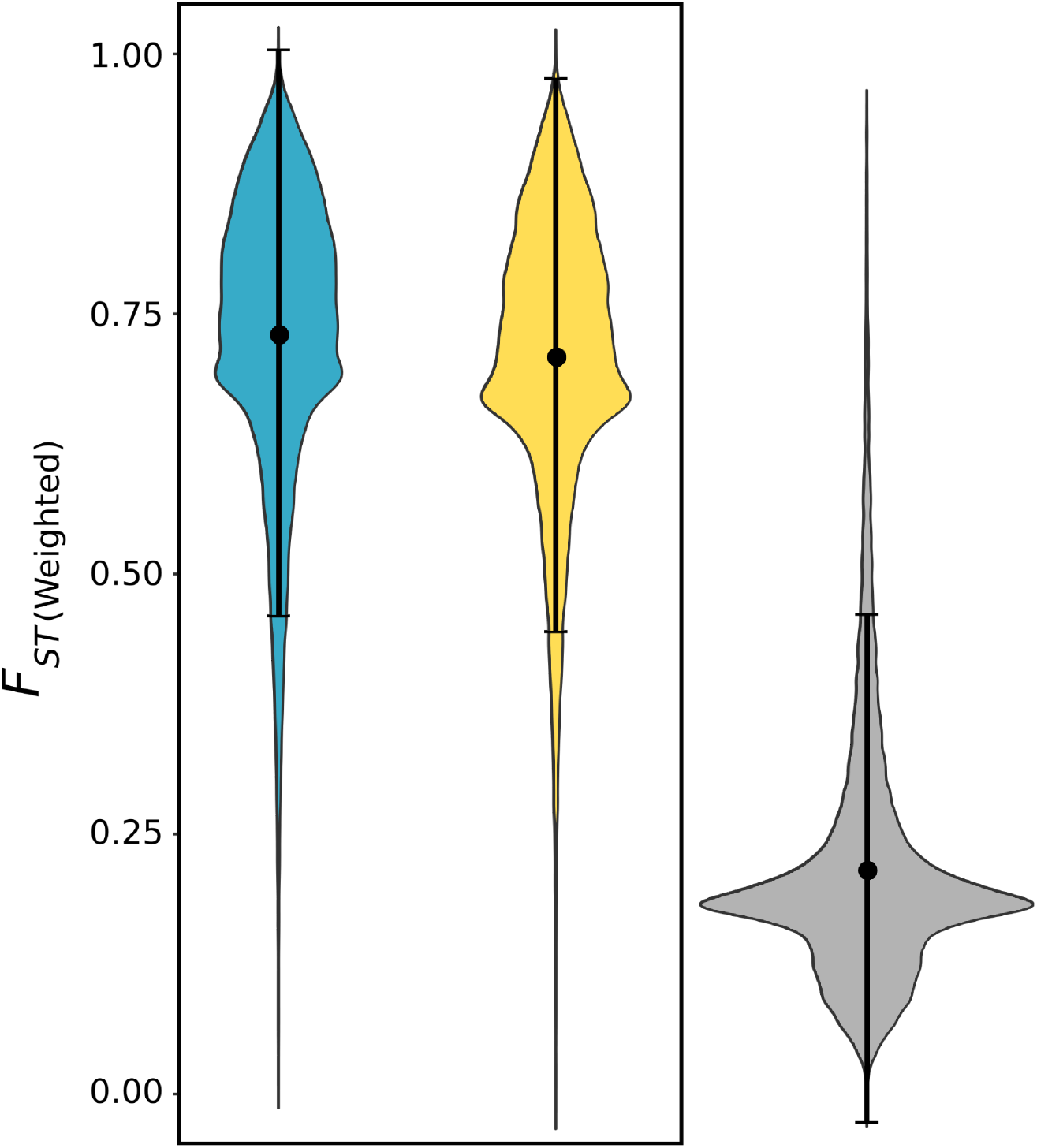
Violin plot of *F*_*ST*_ between Australian *Themeda triandra* clades. *F*_*ST*_ values were calculated across the genome in 50kb windows (10kb slide), and the man value and standard are shown. The first two plots in the box represent the coastal ecotype against the claypan (blue) and inland (yellow) form respectively. The grey plot represents the inland ecotype compared to the claypan form.

